# Lipid peroxidation does not mediate muscle atrophy induced by PSD deficiency

**DOI:** 10.1101/2023.12.22.573082

**Authors:** Hiroaki Eshima, Jordan M. Johnson, Katsuhiko Funai

## Abstract

Mechanisms by which disuse promotes skeletal muscle atrophy is not well understood. We previously demonstrated that disuse reduces the abundance of mitochondrial phosphatidylethanolamine (PE) in skeletal muscle. Deletion of phosphatidylserine decarboxylase (PSD), an enzyme that generates mitochondrial PE, was sufficient to promote muscle atrophy. In this study, we tested the hypothesis that muscle atrophy induced by PSD deletion is driven by an accumulation of lipid hydroperoxides (LOOH). Mice with muscle-specific knockout of PSD (PSD-MKO) were crossed with glutathione peroxidase 4 (GPx4) transgenic mice (GPx4Tg) to suppress the accumulation of LOOH. However, PSD-MKO x GPx4Tg mice and PSD-MKO mice demonstrated equally robust loss of muscle mass. These results suggest that muscle atrophy induced by PSD deficiency is not driven by the accumulation of LOOH.

## Introduction

Accumulation of reactive oxygen species (ROS) has been implicated in the loss of muscle mass induced by physical inactivity [1]. However, targeting ROS to mitigate atrophy has been unsuccessful in human trials. ROS refers to a wide variety of radical molecules whose cellular signals are vast, so it is likely that global suppression of ROS interferes with their other essential roles in biology. Earlier this year, we demonstrated the role of lipid hydroperoxides (LOOH), a class of lipid ROS, in disuse-induced skeletal muscle atrophy [2]. Suppression of muscle LOOH by overexpression of glutathione peroxidase 4 (GPx4) [3], a phospholipid peroxidase, was sufficient to partly rescue the loss of muscle mass induced by hindlimb unloading. Other findings from are consistent with the notion that lipid ROS may mediate the skeletal muscle atrophy induced by physical activity [4-7].

Mitochondria are important sources of cellular ROS, and disuse is known to increase the production of mitochondrial ROS in skeletal muscle. As one potential mechanism for increased ROS, levels of mitochondrial membrane phosphatidylethanolamine (PE) is reduced with disuse [8]. Reduction in the level of mitochondrial PE by deletion of phosphatidylserine decarboxylase (PSD) [9], an enzyme that generates mitochondrial PE, promotes oxidative stress and muscle atrophy. These findings suggest that low level of mitochondrial PE may mediate ROS-induced muscle atrophy with physical inactivity.

The identity of ROS induced by PSD deletion that promotes atrophy remains elusive. Overexpression of mitochondrial-targeted catalase (mCAT) [10, 11] successfully scavenges matrix-resident hydrogen peroxides, one of the main ROS products of the electron transport chain, but is insufficient to rescue muscle atrophy induced by PSD deletion [8]. Similarly, mCAT overexpression does not protect against muscle atrophy induced by hindlimb unloading [12]. In this study, we set out to test whether LOOH scavenging would be protective of muscle atrophy induced by PSD deletion. Because GPx4 overexpression was protective of muscle atrophy induced by hindlimb unloading, we hypothesized that GPx4 would exhibit similarly protective effect on wasting on PSD-deficient muscles.

## Methods

### Animal models

Conditional PSD-knockout (PSDcKO+/+) mice were crossed with tamoxifen-inducible, skeletal muscle–specific Cre recombinase (HSA-MerCreMer+/-) [13] mice to generate PSDcKO+/+; HSAMerCreMer-/- (control) and PSDcKO+/+; HSA MerCreMer+/- (PSD-MKO) mice [8]. Transgenic mice overexpressing GPx4 (GPx4Tg+/-) mice were generated previously [3] and crossed with PSD-MKO mice to generate the PSD-MKO x GPx4Tg (GPx4Tg+/−, PSDcKO+/+, HSA−MerCreMer+/−), PSD-MKO, GPx4Tg (GPx4Tg+/−, PSDcKO+/+, no Cre), and control (PSDcKO+/+, no Cre) mice. PSD-MKO x mCAT mice were previously described [8]. Genotypes were determined via PCR. Cre control mice (PSDcKO−/−, HSA-MerCreMer+/−) and tamoxifen-untreated control mice displayed no difference in phenotype to loxP control mice. All mice were bred onto C57BL/6J background and were born at normal Mendelian ratios. Mice were maintained on a 12 hr light/12 hr dark cycle in a temperature-controlled room. All mice underwent intraperitoneal injection of tamoxifen for 5 consecutive days (7.5 μg/g body mass). Mice were fasted for 4 hours prior to anesthetization via intraperitoneal injection of 80 mg/kg ketamine and 10 mg/kg xylazine and tissue harvest. The tibialis anterior (TA), extensor digitorum longus (EDL), soleus (SOL), plantaris (PLA), gastrocnemius (GAS), and quadriceps (QUAD) were weighed shortly after dissection. All experimental procedures were approved by the University of Utah Institutional Animal Care and Use Committee.

### Western blot

Whole muscle lysate was utilized for western blotting. Frozen GAS muscle was homogenized in a glass homogenization tube using a mechanical pestle grinder with homogenization buffer (50 mM Tris pH 7.6 5 mM EDTA 150 mM NaCl 0.1% SDS 0.1% sodium deoxycholate 1% triton X-100 protease inhibitor cocktail). After homogenization, samples were centrifuged for 15 min at 12,000 X g. Protein concentration of supernatant was then determined using a BCA protein Assay Kit (Thermo Scientific). Equal protein was then mixed with Laemmeli sample buffer and loaded onto 4-15% gradient gels (Bio-Rad). Transfer of proteins occurred on a nitrocellulose membrane which was then Ponceau S. stained and imaged for equal protein loading and transfer quality. Membranes were then blocked for 1 hr. at room temperature with 5% bovine serum albumin in Tris-buffered saline with 0.1% Tween 20 (TBST). The membranes were then incubated with primary antibody for 4-hydroxynonenal (4-HNE; ab48506; Abcam) and GPx4 (ab125066; Abcam). Following incubation, blots were washed in TBST, incubated in appropriate secondary antibodies (Anti-rabbit IgG, 7074, Cell Signaling Technology; or anti-mouse IgG, 31450, Thermo Scientific), and washed in TBST. Membranes were imaged utilizing Western Lightning Plus-ECL (PerkinElmer) and a FluorChem E Imager (Protein Simple).

### Statistical Analyses

Values are presented as means ± SEM. Analyses were performed using GraphPad Prism 10.1.0 software. Statistical comparisons were made using a two-way analysis of variance (ANOVA) and were followed by the appropriate multiple-comparison test. For all tests, P-value less than 0.05 was considered statistically significant.

## Result

We previously showed that mCAT overexpression did not rescue the lethality and muscle atrophy induced by PSD deletion. However, mCAT overexpression did not rescue accumulation of 4-hydroxnonenal (4-HNE) in skeletal muscle of PSD-MKO mice (Fig. 1A). In this study, we crossed the PSD-MKO mice with mice that overexpressed GPx4 (Fig. 1B). This strategy successfully yielded PSD-MKO x GPx4 mice whose muscle HNE was suppressed (Fig. 1C).

**Figure 1.**
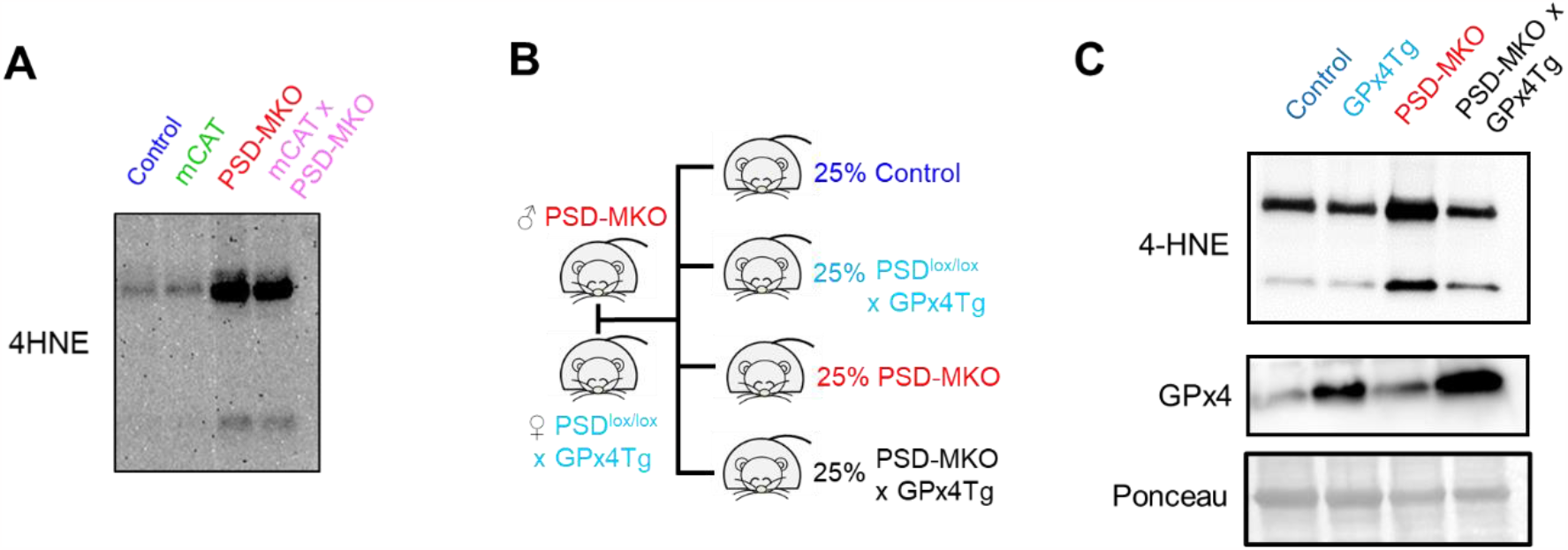
Overexpression of GPx4 rescues the accumulation of LOOH induced by PSD deficiency. (A) Representative 4-hydroxynonenal (4-HNE) western blot image from that PSD-MKO mice that were crossed with mCAT transgenic mice. (B) A breeding schematic to cross PSD-MKO mice with GPx4 transgenic mice to generate PSD-MKO x GPx4 mice. (C) Representative 4-HNE and GPx4 western blot and Ponceau S images from PSD-MKO x GPx4 mice and other groups.

Based on our previous observations that suppression of LOOH/4-HNE is effective in ameliorating disuse-induced muscle atrophy, we hypothesized that LOOH scavenging by GPx4 overexpression would protect these mice from loss of body weight and muscle mass induced by muscle-specific PSD deletion. However, GPx4 overexpression did not rescue decreased body weight induced by PSD deletion (Fig. 2) and were equally susceptible to lethality caused by respiratory failure. Consistent with these data, GPx4 overexpression did not ameliorate atrophy of hindlimb muscle induced by PSD deletion (Fig. 3A-G). Thus, neutralization of LOOH with overexpression of GPx4 was not protective against muscle atrophy induced by PSD deficiency.

**Figure 2.**
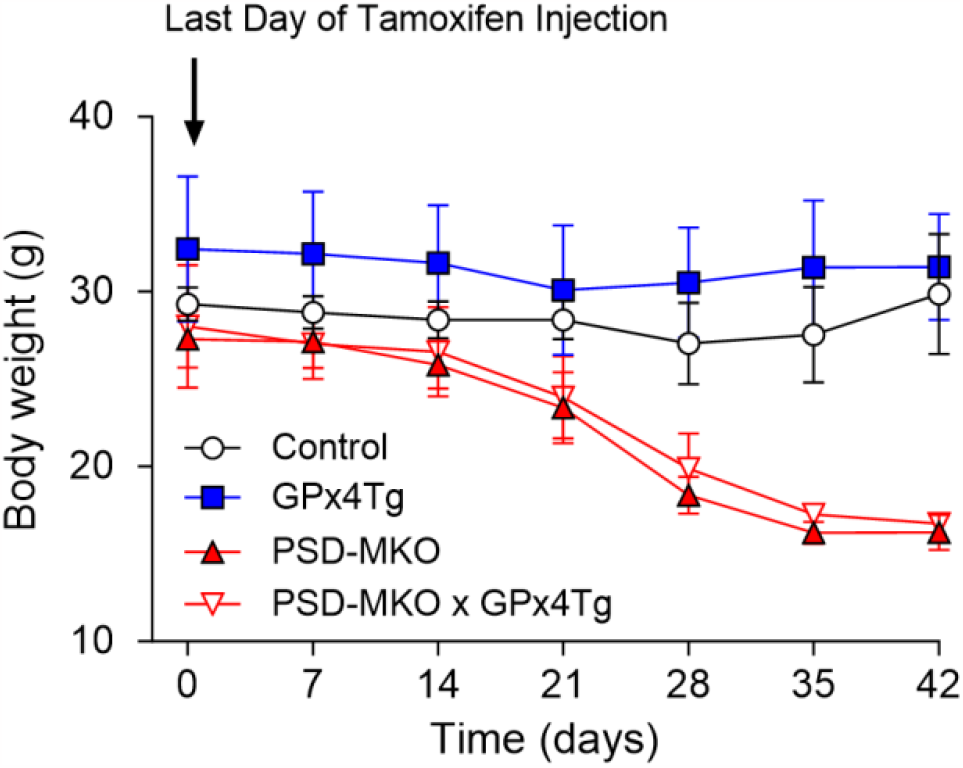
Overexpression of GPx4 does not rescue the loss of body weight induced by PSD deficiency. Body weights after tamoxifen injection (Control: *n* = 5, GPx4Tg: *n* = 3, PSD-MKO: *n* = 4, PSD-MKO x GPx4Tg: *n* = 2).

**Figure 3.**
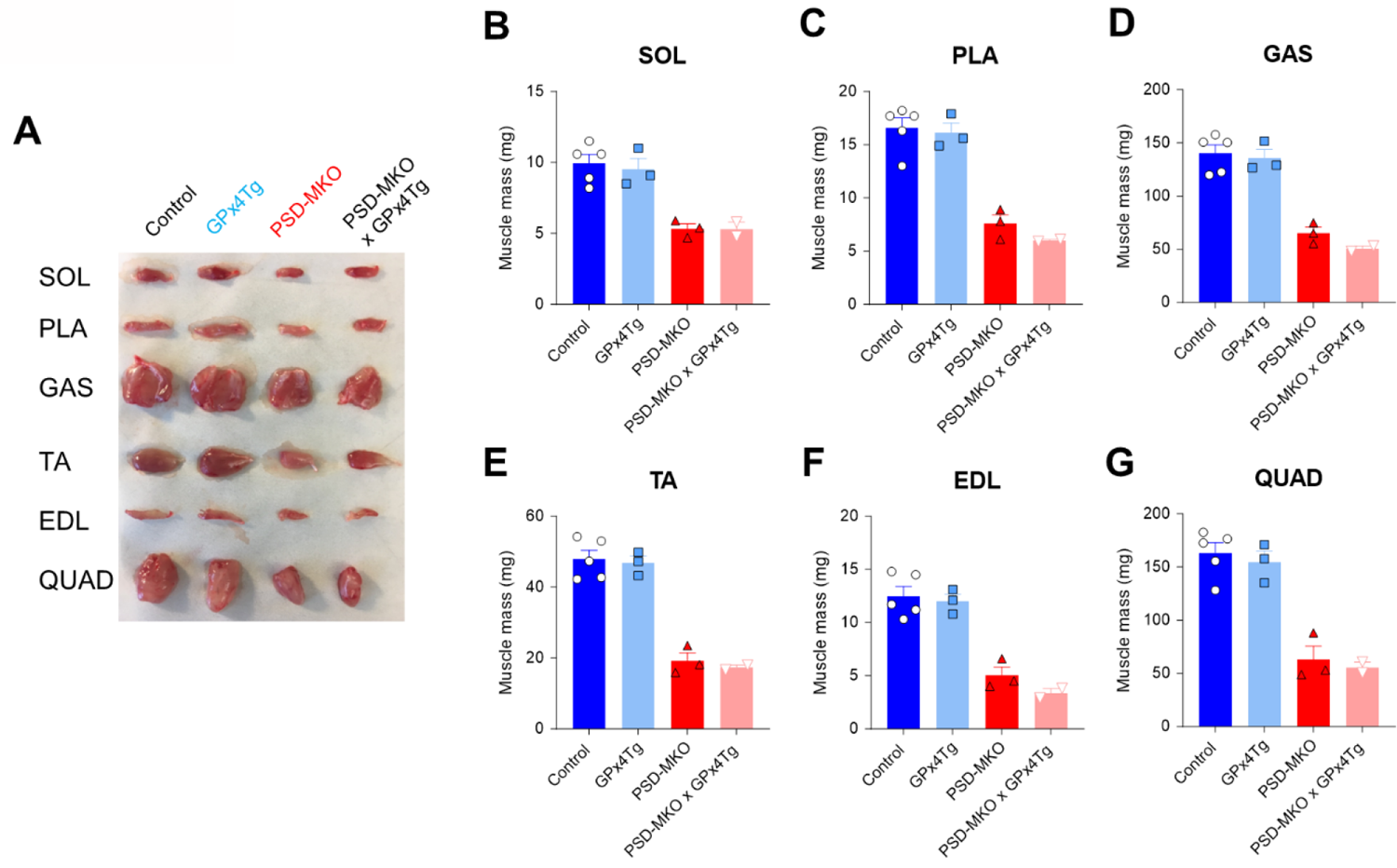
Overexpression of GPx4 does not rescue muscle atrophy induced by PSD deficiency. (A) Representative appearances of soleus (SOL), plantaris (PLA), gastrocnemius (GAS), tibialis anterior (TA), extensor digitorum longus (EDL), and quadriceps (QUAD) muscles. (B) SOL, (C) PLA, (D) GAS, (E) TA, (F) EDL, and (G) QUAD muscle masses (Control: *n* = 5, GPx4Tg: *n* = 3, PSD-MKO: *n* = 3, PSD-MKO x GPx4Tg: *n* = 2).

## Discussion

The PSD enzyme is localized in the IMM and converts phosphatidylserine (PS) to phosphatidylethanolamine (PE) [14], a molecule that likely plays important roles in mitochondrial homeostasis including bioenergetics. The mechanisms by which PSD deficiency promotes muscle atrophy is unclear. Loss of mitochondrial PE leads to electron leak and oxidative stress [8, 15]. However, neutralization of matrix-resident hydrogen peroxide is not sufficient to ameliorate the phenotype induced by loss of PSD [8]. In this study, based on our observation that mCAT overexpression does not rescue the increase in LOOH-induced by PSD deficiency, we examined whether GPx4 expression would be effective in neutralizing LOOH and protecting muscle loss induced by PSD deletion. Our findings unequivocally show that GPx4 expression has no effect on muscle wasting, suggesting that PSD promotes muscle atrophy in an LOOH-independent manner.

LOOH is implicated in ferroptosis, a non-apoptotic form of cell death that is dependent on oxidized phospholipids. Disuse promotes an accumulation of LOOH in skeletal muscle [2, 6]. Global or muscle-specific suppression of LOOH was sufficient to prevent accumulation of LOOH in skeletal muscle and protected muscles from disuse-induced muscle atrophy [2, 4-7]. Together with findings from the current study that GPx4 overexpression does not rescue muscle atrophy in PSD-MKO mice, we can interpret that LOOH drives muscle atrophy with hindlimb unloading but not with PSD deficiency. It remains possible that disuse promotes atrophy by parallel mechanisms mediated by LOOH and reduced mitochondrial PE. This can be tested by performing hindlimb unloading on mice with muscle-specific overexpression of PSD [8].

Recent reports describe that mutations in the human *PISD* gene (that encodes the PSD enzyme) also promotes short stature, in addition to congenital cataracts, facial dysmorphism, platyspondyly, ataxia, and/or intellectual disability [16-18]. Our previous findings that muscle-specific PSD deletion promotes atrophy is consistent with the short stature in these individuals. The condition is also characterized by mitochondrial dysfunction, that is ameliorated by ethanolamine treatment in vitro [16], consistent with the notion that loss of PE contributes to the disease. Our current studies suggest that accumulation of LOOH is not the cause of short stature in these individuals.

In conclusion, PSD deficiency causes muscle atrophy that is concomitant to an increase in LOOH. Even though LOOH has been implicated in muscle atrophy, LOOH induced by PSD deficiency does not appear to mediate its muscle wasting. It remains possible that PSD deficiency contributes to disuse-induced muscle atrophy in an LOOH-independent manner. Maintenance of skeletal muscle mass and function is an energetically expensive process, so it is reasonable to speculate that disuse triggers multiple, some redundant, mechanisms to remove unused biomass and force-generating capacity from the muscle. The study also highlights that the short stature in human PISD deficiency is likely not mediated by LOOH in skeletal muscle and potentially in other tissues.

## Acknowledgements

This research is supported by NIH grants AG074535, DK127979, DK107397, and GM144513 (to K.F.), American Heart Association grant 19PRE34380991 (to J.M.J.), and Uehara Memorial Foundation (to H.E.).

